# The power of a multivariate approach to genome-wide association studies: an example with ***Drosophila melanogaster*** wing shape

**DOI:** 10.1101/108308

**Authors:** William Pitchers, Jessica Nye, Eladio J. Márquez, Alycia Kowalski, Ian Dworkin, David Houle

## Abstract

Due to the complexity of genotype-phenotype relationships, simultaneous analyses of genomic associations with multiple traits will be more powerful and more informative than a series of univariate analyses. In most cases, however, studies of genotype-phenotype relationships have analyzed only one trait at a time, even as the rapid advances in molecular tools have expanded our view of the genotype to include whole genomes. Here, we report the results of a fully integrated multivariate genome-wide association analysis of the shape of the *Drosophila melanogaster* wing in the Drosophila Genetic Reference Panel. Genotypic effects on wing shape were highly correlated between two different labs. We found 2,396 significant SNPs using a 5% FDR cutoff in the multivariate analyses, but just 4 significant SNPs in univariate analyses of scores on the first 20 principal component axes. A key advantage of multivariate analysis is that the direction of the estimated phenotypic effect is much more informative than a univariate one. Exploiting this feature, we show that the directions of effects were on average replicable in an unrelated panel of inbred lines. Effects of knockdowns of genes implicated in the initial screen were on average more similar than expected under a null model. Association studies that take a phenomic approach in considering many traits simultaneously are an important complement to the power of genomics. Multivariate analyses of such data are more powerful, more informative, and allow the unbiased study of pleiotropy.

## Introduction

Forward genetic analyses are generally built on a single measurable quantity, such as size, color, or the presence/absence of a distinctive organismal feature. The rise of phenomics, with its emphasis on high-throughput measurement of high-dimensional traits, is beginning to allow us to address the genetics of more complex traits that no single measurement can capture (Houle 2010; Houle *et al.* 2010). For instance, any one measurement of the wing of a fly, such as the length, incompletely captures wing size and shape (Mezey and Houle 2005; Houle and Fierst 2013; Pitchers *et al.* 2013).

Despite the growing enthusiasm for a more comprehensive approach to the phenotype, the vast majority of genome-wide association studies (GWAS) that include more than one trait have undertaken multiple univariate analyses for each site, rather than a single multivariate analysis (e.g., Teslovich *et al.* 2010; Battle *et al.* 2014). Statisticians have long appreciated the value of genuinely multivariate approaches to association studies (Lange *et al.* 2003; Shriner 2012), and this has led to a recent flowering of multivariate methods and software (O’Reilly *et al.* 2012; Stephens 2013; van der Sluis *et al.* 2013; Scutari *et al.* 2014; Zhou and Stephens 2014; Schaid *et al.* 2016). While these methods are diverse, a consistent result is that multivariate analyses increase the power to detect associations, and the biological usefulness of the results. Given these advantages, it is unfortunate that just a few genuinely multivariate empirical association studies have been published (e.g., Anderson *et al.* 2011; Topp *et al.* 2013). The majority of published multivariate analyses have been examples in the method development papers.

To better understand how a multivariate analysis can result in more biologically interpretable results, consider a typical univariate GWAS in which a single nucleotide polymorphism (SNP) is detected that affects a quantitative trait; for example, a minor allele that increases the length of the *Drosophila* wing. Replicating this effect using an independent sample is the typical next step. However, if the sample size is large enough, the statistical power to detect an effect may be high. Thus, verification will depend on whether the direction of the effect is the same, in this example whether the minor allele also increases wing size in the new sample. There is a 50% chance that a significant effect will be in the same direction by chance alone, making apparent confirmation relatively likely even in the absence of a genuine effect. With multivariate data, however, the original analysis estimates a vector of effects on all the studied traits simultaneously. Confirmation then requires that new estimates replicate the relative magnitude of the effects, and therefore the direction of effects in phenotype space. If large numbers of traits are measured, this is very unlikely to occur in the absence of a biological effect in that direction.

Over the past forty years, evolutionary biologists and quantitative geneticists have developed tools and theory constructing fundamentally multivariate approaches to assessing the genetic basis of phenotypic variation, capturing both the shared and unique attributes of these “traits” (Lande 1979; Hansen and Houle 2008; McGuigan and Blows 2010). Such approaches have proved particularly fruitful in addressing questions about evolutionary diversification (Langerhans and DeWitt 2004), and the response to natural and artificial selection (Blows 2002; McGuigan *et al.* 2005; Hunt *et al.* 2007; Hine *et al.* 2011). Similar advances have occurred in the measurement of phenotypes. For example, the tools of geometric morphometrics and related methods allow us to synthesize comprehensive measures of organismal form (Zelditch *et al.* 2004). Analyses of these data can then disentangle the influence of genotypic effects on size and shape (Weber *et al.* 1999; Weber *et al.* 2001; Palsson *et al.* 2004; Weber *et al.* 2005; Dworkin and Gibson 2006; Klingenberg 2010).

In this paper, we apply a fully integrated multivariate analysis to a genome-wide association data in *Drosophila melanogaster,* drawing on genotypes in the Drosophila Genome Reference Panel (DGRP) (Mackay *et al.* 2012). We analyze the genetic architecture of segregating variation for a 58-dimensional representation of wing shape in (Figure 1A), a model complex trait. We then experimentally validate associations using both an independent panel of inbred lines with targeted genotyping, and using RNAi mediated gene knockdown to examine the degree of replicability for direction of phenotypic effects.

**Figure 1.**
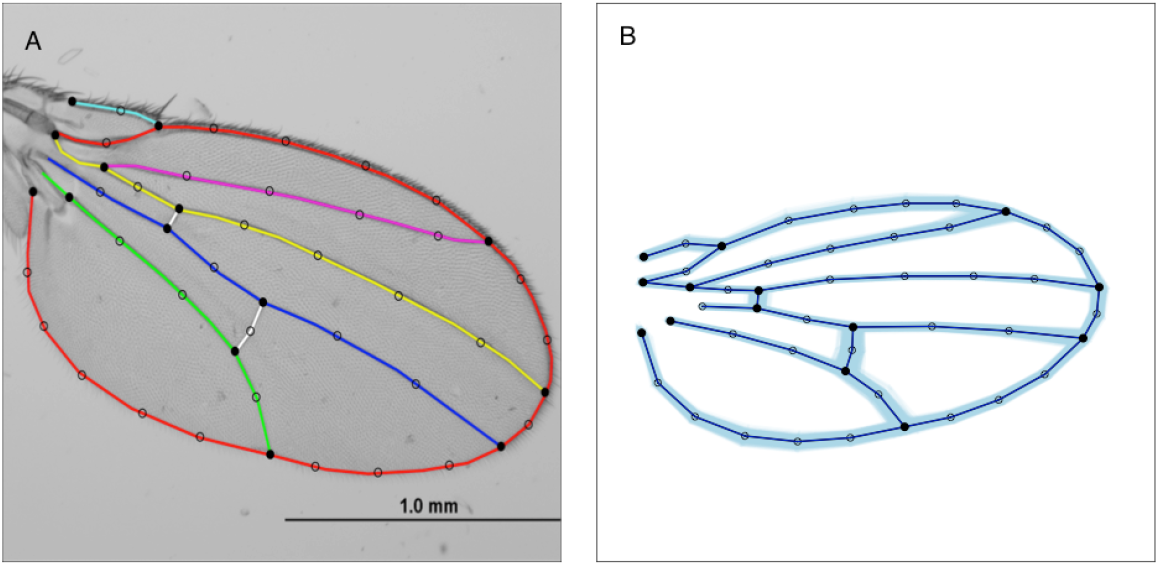
Capturing wing shape with a spline model. Closed circles are landmarks formed by the intersection of splined veins, open circles semi-landmarks used to represent the locations of veins. (A) Splines fit to a typical *Drosophila melanogaster* wing. Colored lines are the splines. (B) Blue overlay represents the range of shape variation among DGRP lines.

## Material and Methods

### Drosophila strains

For the genome wide association study, we used the “Drosophila Genome Reference Panel” (DGRP), a set of inbred lines established from iso-female lines collected at a farmers market in Raleigh, North Carolina (Mackay *et al.* 2012). We obtained phenotypic data from 184 lines scored in Freeze 2 of the DGRP genotyping (Huang *et al.* 2014).

### Rearing, handling of flies and imaging of wings

Wings of DGRP flies were phenotyped independently in both the Houle lab in Florida and Dworkin lab in Michigan. In both labs, each line was reared in vials for at least one generation in the experimental conditions prior to the start of the experiment.

In the Houle lab, flies were reared in a series of 10 temporal blocks over a 14 month period. Twenty-four lines were reared and measured in two or more blocks. Experimental flies were reared in 6 dram vials on a corn meal-sucrose medium preserved with propionic acid, no live yeast added, at 25°C and 55% relative humidity. Each vial was initiated with four parental males and females per vial, who were allowed to lay eggs for three to four days until visual inspection suggested that a sufficient number of larvae had been obtained. The parents were discarded, and the experimental progeny were transferred to new vials of no more than 20 adults to avoid wing damage due to overcrowding. The dorsal surface of the left wings of live flies were imaged using the ‘Wingmachine’ system (Houle *et al.* 2003) using Optem macroscopes with an integrated camera. Annotation, scale information, images as grey-scale TIFF files and guide landmarks were recorded using Image-Pro Plus software (Versions 4, 5 and 6). We sought to obtain images from at least 40 flies per line (20 of each sex). After excluding damaged wings and unsplinable images, data was obtained for a total of 7878 wings from 182 lines, for a mean of 43.3 wings per line. We obtained data from fewer than 40 wings in 23 lines, and from less than 30 wings for only four lines.

In the Dworkin lab, flies were reared in an incomplete balanced block design. Experimental flies were reared in bottles on a cornmeal-molasses-yeast-based medium with carageenan as a gelling agent and propionic acid and methyl paraben as preservatives. Blocks consisted of two replicate bottles of each line reared using food made from the same batch. Each block contained lines that had been reared previously for comparison. Media was physically scored and live yeast was added prior to introducing adult flies to promote egg-laying. Flies were reared separately at 24°C, 60% relative humidity at low density (10 pairs of adult flies per bottle) in a Percival incubator. After 3-5 days (depending on egg density) adults were transferred to new bottles. While eggs were not counted, density was controlled for qualitatively, by removing adults once the desired low egg density was approximately achieved. For those lines with low fecundity, adults were left a few days longer (up to 7 days). After 3-5 days in the second bottle, adult flies were discarded. Water, yeast and paper towel were added to bottles as needed to provide an optimal environment for the larvae. After eclosion and hardening of the cuticle, flies were stored in 70% ethanol at room temperature prior to dissection. Bottles were checked daily as needed until a sufficient number of flies was collected. We dissected between 20–24 wings (left wing of each fly) for each replicate/sex/line.

Wings were imaged at 40X magnification using an Olympus DP30BW camera mounted on an Olympus BX51 microscope and controlled with DP controller software V3.1.1. Images were saved in greyscale as TIFF files. We used the program ‘tpsDig2’ (Rohlf 2011) to record annotation and the guide landmarks. After excluding damaged wings or unsplinable images, data was obtained for a total of 16,272 wings from 165 lines, for a mean number of wings/line of 98.6. We obtained data from fewer than 40 wings in 9 lines, and from less than 30 wings for only four lines.

In total, we obtained phenotypic data from 24,672 wings from 184 DGRP lines, with an average sample size of 134.1 wings/line. One-hundred and sixty-three lines were measured in both labs. We obtained a total of less than 40 measurements (minimum 15) for only four lines.

## Morphometric data

To capture landmark and semi-landmark data from the recorded images, we followed a modified protocol from (Houle *et al.* 2003). Splining and error correction was accomplished in the Java program Wings 3.72 (Van der Linde 2004-2014). Wings fits nine cubic B-spline functions to the veins and margins of wings in the saved TIFF images (Figure 1A), using the locations of the two starting guide points to initiate fitting. Images with outlier splines were reexamined in Wings 3.72, and corrected using a visual editing function if necessary.

The program CPR (Márquez 2012-2014) was used to extract 14 landmark and 34 semi-landmark positions from the fitted splines (as shown in Figure 1). The combined data from the DGRP and validation data sets (a total of 66,890 wings, see below) was subjected to generalized Procrustes superimposition (Rohlf and Slice 1990), which scales forms to the same size, translates their centroids to the same location, and rotates them to minimize the squared deviations around each point. This separates the useful size and shape information from the nuisance parameters introduced by the arbitrary location and rotation of the specimens within the images. The positions of the semi-landmarks were slid along each wing vein (or margin) segment to minimize deviation along the segment. To put numerical results on a more convenient scale we multiplied shape (Procrustes) coordinates by 100.

The 96 superimposed *x* and *y* coordinates from the 48 points recorded generate less than 96 dimensional data, for two reasons. First, each semi-landmark is constrained to lie on a 1 dimensional function, so contributes only 1 degree of freedom (df) to the data. Second, Procrustes superimposition uses 3 df for rotation and translation, and transfers size to a new 1 df variable, centroid size. A 58=2 X 48 – (4+34) dimensional space thus captures all shape variation. The shape data was projected into a 58-dimensional space using principal components analysis of the combined DGRP and validation data, with no adjustment for the fixed sex and lab effects. Thus, PC1 has a large contribution of variation due to the effects of sex. The scores on the first 58 eigenvectors, plus ln centroid size were used for subsequent analyses.

Outliers for the superimposed data were detected in CPR, and then re-examined in Wings 3.72 to allow us to determine whether they represented an unusual wing, or mis-splined specimens, which were corrected. Occasionally a very unusual wing was removed from the data set as an outlier. In all cases, these outlier wings were more than 4 S.D. in Mahalanobis distance from the multivariate mean.

Univariate residuals for shape were generally heavy-tailed (average kurtosis=2.7, defining the kurtosis of a normal distribution as 0). Residuals for principal components 1 and 2 were slightly right-skewed (skew 0.22 and 0.16 respectively), while the remaining shape variables showed no notable skew. Log centroid size was heavy tailed (kurtosis=0.63) and left-skewed (skew=−0.53) Tests for normality of univariate residuals always rejected the normal distribution, which is expected given the large sample size. Association analyses were done on lab, sex and block means (see below), so these departures from normality should have no effect on our results.

## Genetic variation for wing shape

We estimated the genetic variance-covariance matrix due to DGRP line effects using restricted maximum likelihood approach implemented in the mixed model program Wombat (Meyer 2007; Meyer 2010-2015), which we also used to compare the fits of full (40-dimensional) and reduced-rank model likelihoods (Kirkpatrick and Meyer 2004; Meyer and Kirkpatrick 2005; Meyer and Kirkpatrick 2008). Wombat is limited to analysis of 40 variables, so we analyzed the first 39 PCs of shape, plus log centroid size. We fit a pooled sex covariance matrix, treating lab, sex and rearing block as fixed effects. The effect of line was fitted assuming that all lines are equally related. This is an approximation, as 0.05% (11) of all line pairs are more than 50% related (Huang *et al.* 2014).

## DGRP Genotype data

We used the publicly available freeze 2 scoring of line genotypes from February 2013 (Huang *et al.* 2014) (ftp://ftp.hgsc.bcm.edu/DGRP/freeze2_Feb_2013/vcf_files/freeze2.vcf.gz).

Coordinates are based on FlyBaseGenBank Release 5. We used only calls of homozygous genotypes, and treated others as missing data. We refer to all polymorphisms as SNPs, despite the fact that some polymorphisms involve multiple nucleotides. If two or more SNPs were found at the same site, we analyzed the one with the highest minor allele count, treating all rarer variants as equivalent to the reference. We used only homozygous calls with genotypic phred scores ≥20 at sites with exactly two alternative types. All other calls were treated as missing data. Initial analyses to choose sites for validation in the Maine and North Carolina population were carried out using Freeze 1 data from August 2010, available at https://www.hgsc.bcm.edu/arthropods/drosophila-genetic-reference-panel.

## Linkage disequilibrium

Linkage (gametic) disequilibrium (LD) complicates the interpretation of significant associations uncovered in a GWAS. We quantified LD as the squared gametic correlation between sites

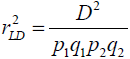

where *p*_1_, *p*_2_, *q*_1_, and *q*_2_ are the major and minor allele frequencies at the two sites, and *D* = *x*_11_ − *p*_1_*q*_1_, where *x*_11_ is the frequency of gametes carrying both the alleles indexed by the frequencies alleles *p*_1_ and *q*_*1*_(Weir 1996). To help us interpret our results, we calculated the gametic correlation between all sites judged to be significant at a false discovery rate (FDR) of 5% (see below) and all other sites throughout the genome and retained a list of all those pairs where the squared correlation *r*^2^>0.5. This cutoff was chosen based on simulations that show that LD of *r*^2^<0.5 will infrequently generate false positives for a SNP in LD with a SNP that has a phenotype effect size typical of those detected in this study in a similar number of lines (Houle and Márquez 2015).

For SNPs judged to be significant in our association tests, we carried out a second analysis designed to separate clusters of SNPs that were highly correlated with many other SNPs from those that were more or less independent. To find an initial set of clusters, we used the SAS/FASTCLUS Procedure (SAS 9.3), which uses *q* vectors of SNP genotypes as seeds to group input SNPs into up to *k* clusters with a radial spread equal to *R*, where *k* and *R* are user-defined parameters. In our clustering algorithm, we first assign a deliberately large value to *k* (=2000), and let FASTCLUS compute optimal seeds. In this step, we also let FASTCLUS automatically impute missing genotype data. In a second run, we submit the previously imputed data to FASTCLUS, and save the output as seeds for subsequent iterations of the same algorithm. We then iterate this step until both the number of clusters and a least squares optimization criterion plateaus. We chose the radius *R* for our clusters to match the *r*^2^>0.5 cutoff. From the law of cosines, the distance, *d*, between two SNP vectors is related to their correlation by 
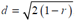
, leading to *R* = 0.7654.

The above algorithm does not ensure that the clusters identified are discrete. To compensate for this we carried out a second, refinement phase. This phase consists of three steps: first, we scan each non-singleton cluster to determine whether any of its members do not conform to the clustering criterion (i.e., its squared correlation with every other member of the cluster does not equal or exceed 0.5). SNPs that violate the criterion are marked as singletons for subsequent processing; second, squared correlations between singleton and all other SNPs are computed to allow for orphan SNPs to join established clusters, or for pairs of singletons to cluster when the *r*^2^ > 0.5 criterion is met. If a SNP is correlated with more than one cluster, it is allowed to join the cluster with the most members; finally, the last step merges clusters with highly correlated SNPs. Specifically, two clusters are combined into a single cluster when the minimum of the maximum squared correlations computed between all pairs of members of different clusters exceeds 0.5. All of these steps were iterated until convergence. The result from our algorithm is a series of clusters comprising SNPs each satisfying the correlation criteria *r*^2^ ≥ 0.5 with every other SNP within the cluster, and *r*^2^ < 0.5 with every SNP that does not belong in the same cluster.

## Impact of chromosomal inversions

Previous inversion karyotyping of the DGRP lines (Corbett-Detig and Hartl 2012; Langley *et al.* 2012; Huang *et al.* 2014) yielded inconsistencies for three inversions that are present in at least seven of the DGRP lines, (In(2L)t, In(2R)NS, In(3R)Mo). Houle and Márquez (2015) found that a principal component analysis on the set of genotypes that had disequilibrium *r*^2^>0.5 with at least 200 other SNPs more than 100kb distant from the focal SNP correctly diagnosed the inversion-type of all but one chromosome arm for which the previous analyses were consistent. Consequently, we used high PC1 genotypic scores as our indicator of karyotype, as detailed in Houle and Márquez (2015).

In lines that were inferred to carry a heterozygous or homozygous alternate karyotype on the basis of the above analysis, we treated all genotype calls as missing in regions in high LD with genotypes typical of the three common alternate karyotypes. For In(2R)NS and In(2L)t this included sites between the breakpoints inferred by Corbett-Detig *et al.* (2012), plus 20kb either side. Corbett-Detig and Hartl (2012) and our own additional analyses suggest that In(3R)Mo is in strong LD with the entire distal part of arm 3R from approximately 16Mb to the distal tip at 32Mb, although the breakpoints of that inversion are approximately 17.2 and 24.9 Mb. Consequently, we masked all sites with coordinates greater than 16Mb in lines inferred to carry an In(3R)Mo genotype.

## Detecting phenotypic associations

The phenotypic data consisted of 58 wing shape variables computed as described above, plus the natural log of centroid size. We thus analyzed wing form in size-shape space (Bookstein 1986; Mardia and Dryden 1989). The data for analysis was mean wing form for each combination of line, lab and sex. We treated genotypes between inversion breakpoints in lines carrying the alternate karyotypes as missing. Our analysis included SNPs where the minor allele was homozygous in at least five lines and where the number of lines called was at least 120 of the 184 freeze2 DGRP lines for which we obtained phenotypic data. This left us with 2,517,547 polymorphic sites.

Effects of SNPs on morphometric variation were quantified using a multivariate linear model taking into account the effects of lab, sex, SNP and line

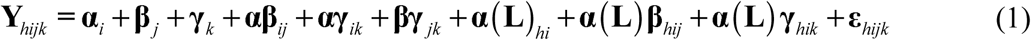

where **α**, **β**, and **γ** represent vectors of fixed effects of the *i*^th^ SNP, *j*^th^ sex, and *k*^th^ lab, respectively, **α**(**L)** represents the random effect of the *h*^th^ inbred line nested within SNP, ε is the residual vector, and higher order terms represent interactions between these factors. To compare the results of multivariate and univariate analyses, we also calculated univariate tests using the model in equation (1).

The above model could not be fit as a mixed model. We approximated the mixed model tests and estimation using the following multi-step procedure. We first estimated the sum of squares and cross-products (SSCP) matrices using a least squares method in SAS Proc GLM, designating terms involving line nested in SNP as random with variates weighted by their sample sizes. Because sample sizes over labs and sexes were always unbalanced, the denominators of within-group SSCP matrices, **W,** were assembled as weighted averages of the SSCP matrices obtained in this first analysis. The weights were obtained from the coefficients of the expected mean squares calculated in a univariate analysis of the same SNP in SAS Proc GLM using the Random/Test option. We assessed the statistical significance of model terms using an *F*-distributed statistic based on Wilks’ Λ (Rao 1973), computed as 
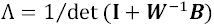
 where **B** is the between-group SSCP matrix.

Least-squares estimates of SNP effect vectors, were obtained from a simpler model neglecting interactions of SNP effects with sex and lab,

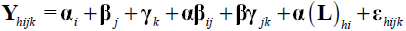

Effect size is the length (norm) of this vector of effects, 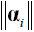. The amount of variance explained is 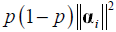, where *p* is minor allele frequency (MAF).

These analyses were written in SAS macros and were run at the High Performance Computing facility at North Carolina State University, the Research Computing Center at Florida State University, and a standalone Linux server at the Biological Science Department at Florida State University.

To control the FDR we applied the Storey and Tibshirani (2003) approach as implemented in the R package fdrtools (Strimmer 2008).

## Gene Ontology Analysis

For each SNP significant at a FDR of 5%, we downloaded the full gene ontology information for the nearest genes upstream and downstream on both the negative and positive strands. We identified the closest coding region to each of these SNPs and used WebGestalt (Wang *et al.* 2013) to test for enrichment of Gene ontology categories using a Benjamini-Hochberg (1995) correction after testing against the null expected distribution of all genes assuming a hyper-geometric distribution.

## Evaluating the influence of correlated SNPs

While it is useful to know that a SNP genotype is significantly associated with a phenotypic effect somewhere in the genome, we also want to determine whether the phenotypic effect is likely to be due to that SNP, or to variants in LD with the SNP. We refer to SNPs (QTNs) that affect the phenotype as causal. To determine which SNPs implicated in the MANOVA analysis were most likely to be causal, we implemented a stepwise multivariate regression algorithm. The goal of this analysis was not to arrive at a best model for SNP effects, but to determine, for each significant focal SNP, how other SNPs in the data set altered the statistical signal from the focal SNP, and to determine where those competitor SNPs map. SNPs whose conditional effects remain significant in models that include other variants are more likely to be causal. SNPs whose conditional significance is diminished by closely linked SNPs, but not by distant variants, may reliably signal the presence of a neighboring QTN.

For each of the significant SNPs we assembled a family of variants to be checked for their influence on the significance of the focal SNP. This family comprised three sets of variants. The first group consists of all significant SNPs that were annotated as being closest to, or within 2kb of the transcript of a gene that the focal SNP is either in, closest to or within 2kb of the focal SNP The second group consists of all SNPs anywhere in the genome that have LD *r*^2^>0.5 with any of the first group of significant SNPs, regardless of whether they are significant when analyzed separately. For SNPs outside the breakpoints of the blocks of LD associated with the three common inversions, we also considered the three common inversion karyotypes (In(2L)t, In(2R)NS, In(3R)Mo). For SNPs between these breakpoints, we performed the multiple regressions for just those lines without evidence of the non-Standard karyotype, as above. Within this universe of possible competitor variants, we screened for SNPs with LD r^2^>0.9, then dropped the SNP with the lowest number of genotype calls in each such pair, while recording the existence of a highly correlated SNP. If the focal SNP mapped between the breakpoints of the blocks of LD associated with the three common inversions, potential competitor SNPs with less than four copies of the minor allele in genotypes scored as the Standard karyotype for that region were dropped from the analysis. The result of this is a family of *t* variants to be examined for their influence on the statistical results for focal SNP *f*. To enable the simultaneous analysis of multiple SNPs, we assigned the common allele to missing calls in all *t* non-focal SNPs and ignored genotypes for lines with missing calls at SNP *f*.

The data for the multivariate regression were the vectors of least-squares means for the shape and size variables from a linear model with lab, sex and DGRP line as main effects, 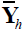. The first step was to fit the regression model 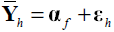, where α_*f*_ is the vector of effects for the focal significant SNP being tested, and record the value of Wilk’s λ and the associated P-value as a baseline for evaluating the change in significance of the focal SNP when additional predictors are used. Next, models of the form 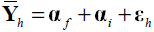, including the focal SNP *f* and one additional SNP, *i*, were fit, yielding *t* round 1 estimates of the significance of SNP *f* when considered in models of with an additional variant. A SNP was listed as a competitor if it was either perfectly correlated with the focal SNP, or reduced the F ratio for the test of the significance by 10% or more, and the ratio of the P value after addition to the P value before addition of the competitor SNP multiplied by the P value from the MANOVA is less than 0.0001.

Finally, we performed up to six stepwise additions of SNPs to this model. If at least one of the *t* variants at round *j* had a P-value<0.001, the variant with the minimum *P* was added to the model. This process was repeated for up to five additional steps, so the final model could have from one (the focal SNP) to seven variants in it. We did not delete SNPs from the model once they entered. We used the results of the round 1 tests to indicate how many variants compete with the focal SNP, and the result from the final model to indicate the overall robustness of the initial MANOVA result. A focal SNP was listed as non-significant in the multiple regression analysis when the ratio of the P value for the final model to the P value before addition of any competitor SNPs multiplied by the P value from the MANOVA is less than 0.0001.

Multivariate regressions were calculated in SAS Proc Reg, using the MTEST statement.

## Quantitative knockdowns of gene activity for validation

We knocked down expression of genes of interest using the Geneswitch Gal4 construct (GS, Roman *et al.* 2001) engineered to be under the regulation of a ubiquitous *tubulin* driver (Tub-5 GS) generously furnished by Scott Pletcher. GS is a chimeric protein that combines yeast Gal4 with a mammalian progesterone receptor. The resulting protein is activated in the presence of a progesterone analog in the diet, which we furnish as the drug mifepristone. There is, however, residual Gal4 activity in the absence of Mifepristone. We backcrossed the Tub-5 GS construct into a wild-type Oregon R (OreR+) background before these experiments. GS was used to drive expression of interfering RNA for a gene of interest (UAS-[GOI] RNAi) constructs obtained from the TRiP project (Ni *et al.* 2008; in a *yv* background) and the Bloomington Drosophila Stock Center or the Vienna Drosophila RNAi Center (Dietzl *et al.* 2007; in a *w*1118 background). The list of RNAi stocks is listed in File S3 3. All knockdown experiments were conducted in the Houle lab.

To carry out a knockdown experiment, we crossed reciprocally crossed Tub-5GS and UAS-[GOI] RNAi stocks, and allowed these flies to lay eggs on media containing at least four different concentrations of mifepristone. We used mifepristone concentrations of 0, 0.3, 0.9, and 2.7 µM in all experiments. Initial experiments include a fifth concentration (0.1 µM), but in most cases there were no phenotypic differences between 0 and 0.1 µM treatments. For each concentration of mifepristone four replicate vials were set up; a fifth replicate was set up for 2.7 µM due to low survivorship in many experiments. We placed ten virgin females with five males in each vial.

Three different control crosses with their respective reciprocals were also set up: Tub-5GS x the appropriate RNAi background (either *yv* or *w*^*1118*
^), UAS-[GOI]RNAi x OreR+, and RNAi background (either *yv* or *w*^*1118*^) x OreR+. Reciprocal and control crosses were set up at the same time on medium from the same batch. After six days, all the parents were moved to fresh vials with the appropriate mifepristone concentration, and then discarded after an additional six days. Offspring were moved to vials with fresh, normal food, sorted by sex, and their wings were imaged at least two days after eclosion. We imaged wings from 20 F_1_ females and males from each treatment in each reciprocal cross.

The distribution of within reciprocal, sex and treatment data was frequently heteroscedastic; higher mifepristone RNAi treatments generally had higher variance, often showing outliers along the major axis of RNAi effects. Consequently, we analyzed the within-sex-treatment-reciprocal medians. Further analyses (in prep.) of control and experimental data suggests that mifepristone has background-specific effects on wing shape across UAS-[GOI]RNAi crosses, and data were adjusted for these effects before further analyses. Finally, we calculated the linear effect of mifepristone on the 58 shape dimensions in a linear model with sex and reciprocal as categorical effects and mifepristone as a continuous predictor. In some cases, the reciprocals differed significantly in their effects, and were analyzed separately. These are designated by the sex of the Tub-5 GS parent in File S3. The parameters of the multivariate regression of mifepristone were retained as the effect vector of the manipulated gene of interest.

## Comparing knockdown vectors to SNP vectors

The correlation of column vectors *x* and *y* is

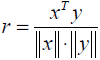

Where ^T^ indicates transpose and 
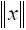
 denotes the length (2-norm) of vector *x*. Like all correlations, − 1 ≤ *r* ≤ 1. In this work, the sign of the correlation is arbitrary, because we could take either the major or the minor allele as the reference, so we report the absolute value of vector correlations. The arc cosine of *r* is the angle between the two vectors. A correlation of 1 means that the vectors point in the same direction, while *r*=0 means that the two vectors are orthogonal (at 90 degrees).

We tested the statistical significance of vector correlations between the knockdown vectors and SNP vectors by comparing the observed correlations to the distribution of correlations under the null hypothesis of no relationship. Even if the SNPs have no real effects, the inferred vectors will tend to fall in the more variable regions of phenotype space, so to ensure that the random vectors were appropriately sampled we took two approaches. First, we assumed that the estimated directions of effects in the overall sample of 2.5 million SNPs were representative of the random distribution of effect directions. Second, we randomly sampled vectors from a multivariate normal distribution with mean 0 and covariance equal to the estimated among-line genetic covariance matrix. These two approaches yielded similar, but not identical, results. We report the results using the random sample of inferred vectors, but both approaches agreed in all the specific cases discussed.

The second major challenge is that many of the significant SNPs are in LD with other SNPs, so their effects will be shared among correlated SNPs. To compensate for these correlated effects, and to minimize the overall number of tests, we computed the average SNP effect over all members of the 862 linkage disequilibrium clusters identified above, weighted by the product of minor and major allele frequencies. Only the 792 clusters where SNPs were less than 5kb apart and where all SNPs were closest to a single gene were tested, as clusters including more distant SNPs are much less likely to implicate the focal gene.

To test for significant correlations, we compared quantiles of vector correlations between the knockdown vector and 10,000 random vectors. When the SNPs nearest one gene were in *k*>1 LD clusters, we generated *k* random correlations in each of the 10,000 replicates and calculated quantiles of these maximal correlation against which to judge the significance of that knockdown.

To judge whether the entire sample of correlations between knockdown and SNP vectors is higher than expected, we calculated samples of 10,000 sets random correlations and calculated the quantiles of the difference in the magnitude of the mean real and random sets of correlations.

To compute an experiment-wise cutoff for which knockdown effects are similar to effects of the 792 single-gene SNP LD, we first calculated 1,000 sets of *k*=792 correlations of random SNP effects with each knockdown vector. The quantiles of the maximum of the 792 correlations were calculated and compared to the observed vector correlations. These quantiles differ considerably with the direction of the knockdown effect; vectors close to the principal axis of genetic variation (PC1) are much more common in the estimated set of vectors, and so have quantiles considerably larger than those in less-common directions. In File S3 we report both quantiles and the vector correlations of each knockdown vector with the first five PCs of the among-line variance matrix.

## Maine and North Carolina (ME-NC) populations for validation

### Flies

Female *D. melanogaster* were collected in the summer of 2004 at a Peach Orchard in West End, North Carolina (NC2), and in a blueberry field in Cherryfield Maine (ME), by Marty Kreitman (Goering *et al.* 2009; Reed *et al.* 2010). All lines were full-sib inbred for 15-20 generations. In total 190 lines were used (~50% from each population). Flies were reared in the Dworkin lab at 25ºC in a 12:12 light/dark cycle at constant 50% humidity, similar to previously described experiments (Dworkin and Gibson 2006). We dissected approximately 20 wings/replicate/line for a total of 7968 male and 7781 female wings from 153 lines. Collectively we refer to this as the ME-NC panel.

### Choice of SNPs to validate

SNPs were chosen for inclusion in the validation panel based on the findings of an analysis using Freeze1 genotypic data. We began by compiling a list of SNPs with the smallest associated *P*-values, removing those where the SNPs effect was unstable (LogRatio of *P* within 1.5 S.D. of 0) and ranking the remainder by effect size.

We then excluded SNPs whose minor allele was present in fewer than 9 DGRP lines, making the assumption that more common SNPs would be more likely to be found in the other populations. SNPs whose associated effect was highly correlated with other SNP effects were also removed from the list to avoid attempting to validate effects associated with SNPs that were in LD with the QTN itself, since patterns of LD may differ among populations. Finally, we retained the 350 SNPs that were closest to the transcript of a gene.

### Genotyping

The genotyping for our validation SNP set was carried out by KBiosciences (Now LGC Genomics) using ‘Kompetitive Allele-Specific PCR’ assays (KASP). This is a fluorescence-based genotyping technology that uses allele-specific primers, making it generally more accurate for smaller jobs than high-throughput methods. We designed primers based on 100 base pairs of the *D. melanogaster* reference genome from Flybase (version 5.41) on either side of each SNP. We submitted these sequence snippets, along with samples of genomic DNA extracted from 15 flies from each of the MENC lines to KBiosciences. Several duplicated control samples (same genotype, but independently labeled) were included to assess any technical variation in genotyping.

### Analysis

For the MENC validation analysis, we used the same pipeline and analysis framework as described above for the DGRP, but excluding the effects of lab and sex (as only males were phenotyped). To determine if the average vector correlation of the true DGRP SNPs with the MENC SNPs was greater than expected by chance, we repeatedly computed the vector correlations between random subsets of DGRP SNPs (350 at a time) with the MENC SNPs, and estimated the average across the set. This process was repeated 1000 times.

## Results

### High repeatability for wing shape across labs

We obtained independent measurements from 163 DGRP lines in both the Dworkin and Houle labs, out of a total of 184 lines phenotyped. The eigenvectors used to score the data and the means and standard deviations of the variables by sex and lab are shown in File S1. Wing shape shows considerable variation among lines (Figure 1B, Figure 2A). As described in the Methods, each lab reared flies slightly differently. The factors that differed include rearing temperature, food, and measurement hardware, as well as unknown aspects of lab routine. Despite these environmental differences, line effects on wing shape have a high degree of inter-lab repeatability with respect to both effect sizes (Figure 2A) and directions (Figure S1A). Wing size was, however, weakly correlated across labs (Figure 2B). This is likely to be due to genotype-environment interactions with lab rearing practices, rather than measurement error, as repeatability of wing size within lab is high (Figure S1B). A MANOVA on line-sex-lab means shows that the effects of lab (Wilk’s λ=0.0026, F=3014.6, df=(59,460), sex (Wilk’s λ=0.027, F=275.9) and lab by sex interactions (Wilk’s λ=0.35, F=14.2) are all highly significant (P<0.0001), reflecting subtle differences in means across labs.

**Figure 2.**
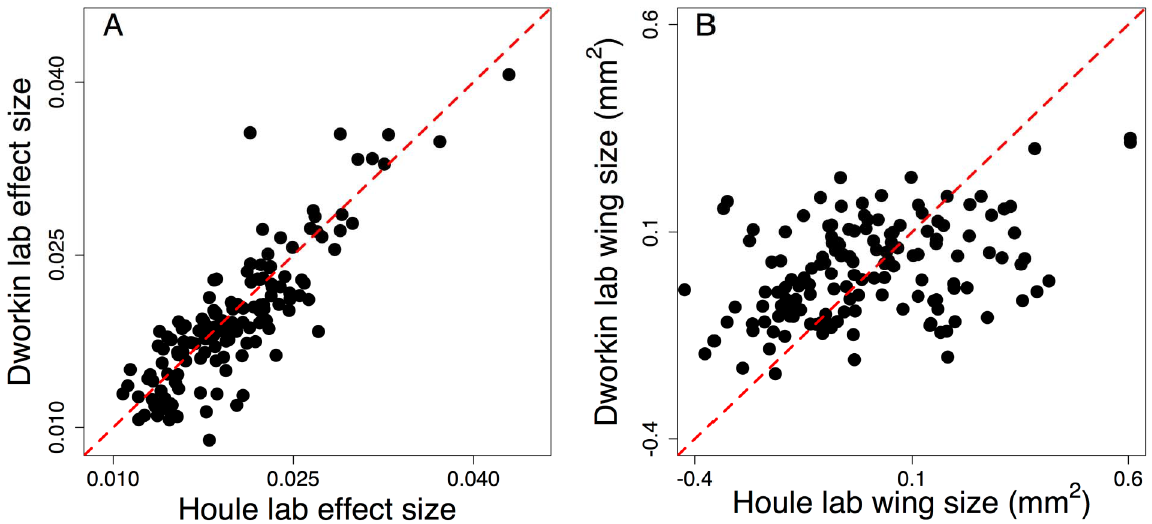
Inter-lab repeatability. A) High repeatability of line effect sizes across labs. B) Low repeatability of size across labs, despite high intra-lab repeatability (S1B Figure).

### Genetic variance in wing shape in the DGRP

To verify the presence of genetic variance in wing size and shape, we estimated the variance-covariance matrices in wing size and shape. Due to software limitations, we could test for genetic variance in only 40 dimensions (out of 59 possible), so we chose to fit the first 39 principal component (PC) scores for wing shape, plus log_10_ centroid size. A model with genetic variance in all 40 possible dimensions fit better than models with 39 dimensions by 1,466 penalized log-likelihood (AICC) units. This is strong evidence that at least 40 independent aspects of wing shape are affected by genotypic variation in the DGRP sample.

### Chromosomal inversions influence wing shape, but Wolbachia does not

Three inversion karyotypes (In(2L)t, In(2R)NS, and In(3R)Mo) were found in more than four of the DGRP lines that we phenotyped (Huang *et al.* 2014; Houle and Márquez 2015). Approximately 50% of the lines carried the intracellular parasite *Wolbachia* (Huang *et al.* 2014). We conducted MANOVAs on the effects of inversion genotypes and *Wolbachia* status, with the results shown in Table 1. Each of the three inversions has a highly significant effect on wing shape-size, but *Wolbachia* infection status has no significant effect.

**Table 1.** **Summary statistics for MANOVAs for the effects of the three common inversions and Wolbachia infection status on wing shape-size.**

|  | numerator | denominator |  |  |
| --- | --- | --- | --- | --- |
| Predictor | df | df | Wilk's $\lambda$ | $P$ |
| In(2L)t | 128 | 94.7 | 0.364 | $2.08 \times 10^{-7}$ |
| In(2R)NS | 128 | 93.4 | 0.475 | $2.70 \times 10^{-3}$ |
| In(3R)Mo | 128 | 93.8 | 0.327 | $4.53 \times 10^{-9}$ |
| Wolbachia | 59 | 124.7 | 0.660 | 0.34 |

### Basic GWAS analysis

After excluding calls in inverted regions in lines identified as carrying one of the three common inversion karyotypes, we were left with 2,517,547 polymorphisms with minor allele count ≥ 5. For convenience, we refer to all polymorphisms as single nucleotide polymorphisms (SNPs), despite the fact that some involved indel variation or multiple nucleotides. We carried out individual MANOVAs of the effect of genotype on wing shape for each SNP. To pick out SNPs for additional analyses, we used the false discovery rate (FDR) algorithm of Storey and Tibshirani (2003), bearing in mind that all such methods assume independence of each analysis. A total of 2,396 sites had significant effects using a 5% FDR cutoff (q-value < 0.05). This analysis estimated that the *P*-values can be explained by mixture of η_0_ = 71.5% SNPs with no phenotypic effect, with the remainder having some effect. Figure 3A shows a Manhattan plot of the multivariate results. A list of the significant sites, test statistics, effect sizes, variance explained, plus information about genes implicated are given in File S2.

**Figure 3.**
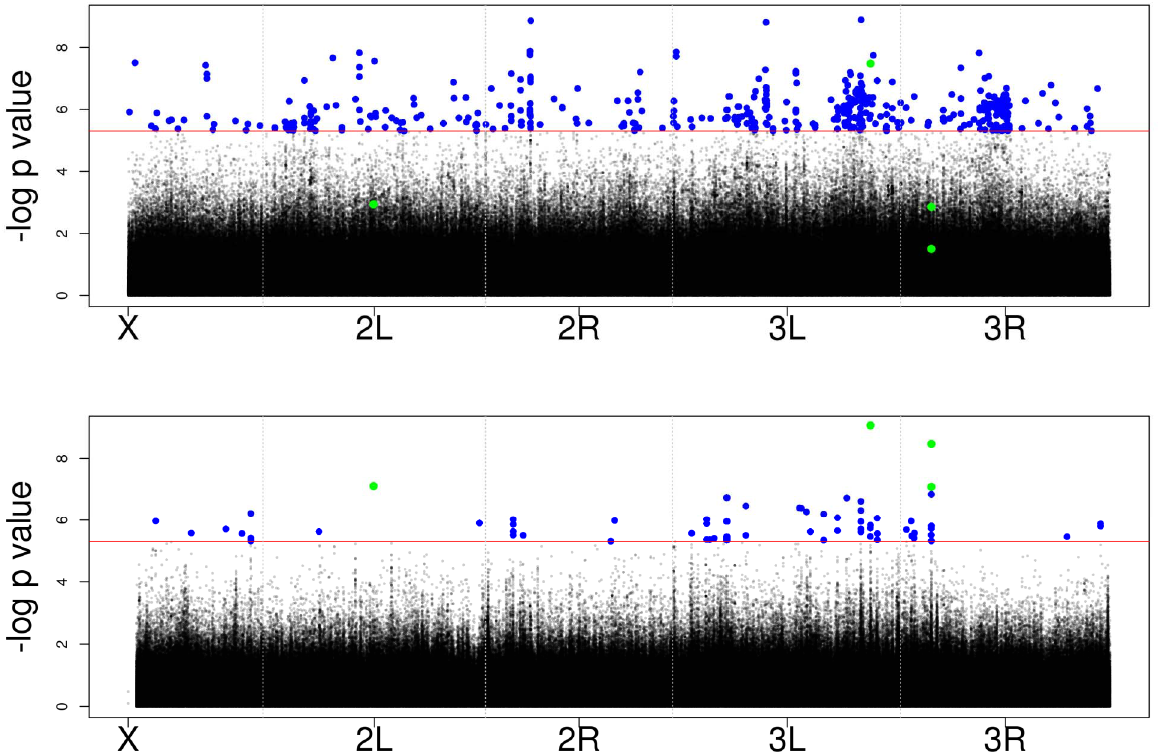
Manhattan plots of the log_10_ inverse P-values from (A) multivariate analysis and (B) a univariate analysis of PC1. Solid red line is *P*=0.00007, the cutoff for a 5% FDR using the Storey and Tibshirani analysis of the multivariate data. Green points are the four SNPs that reach the 5% FDR cutoff from analysis of just the PC1 *P*-values.

We calculated the genetic variance in shape-size explained by each of the significant SNPs as a proportion of the trace of the among-line genetic variance-covariance matrix. Estimated effect sizes are modest, and no single SNP is estimated to explain more than 3.6% of the variance. In addition, the estimated effect sizes are clearly too large on average, as the mean percentage of variance explained is 1.4% (median is 1.3%). These results are consistent whether considering the shape-only data or shape and size simultaneously, which are almost perfectly correlated (0.99). There are two known causes for the upwards bias in effect size. First, sampling variation causes effect for SNPs judged to be significant to be overestimated (Beavis 1994; Beavis 1998; Xu 2003). Second, these analyses do not compensate for the effects of linkage disequilibrium, which we return to below.

A quantile-quantile plot of the *P* values is shown in Figure 4. For sites with minor allele frequency (MAF)<0.15, the distribution shows clear evidence of substantial deviation from the expected uniform distribution throughout the range of P values. We interpret this as largely due to the spreading signal of true effects to the large number of sites in LD with rare alleles (see below). The *P* values are much closer to the null distribution at sites with MAF>0.15. This distribution is also consistent with a very large number of sites each having small phenotypic effects. We return to these issues below.

**Figure 4.**
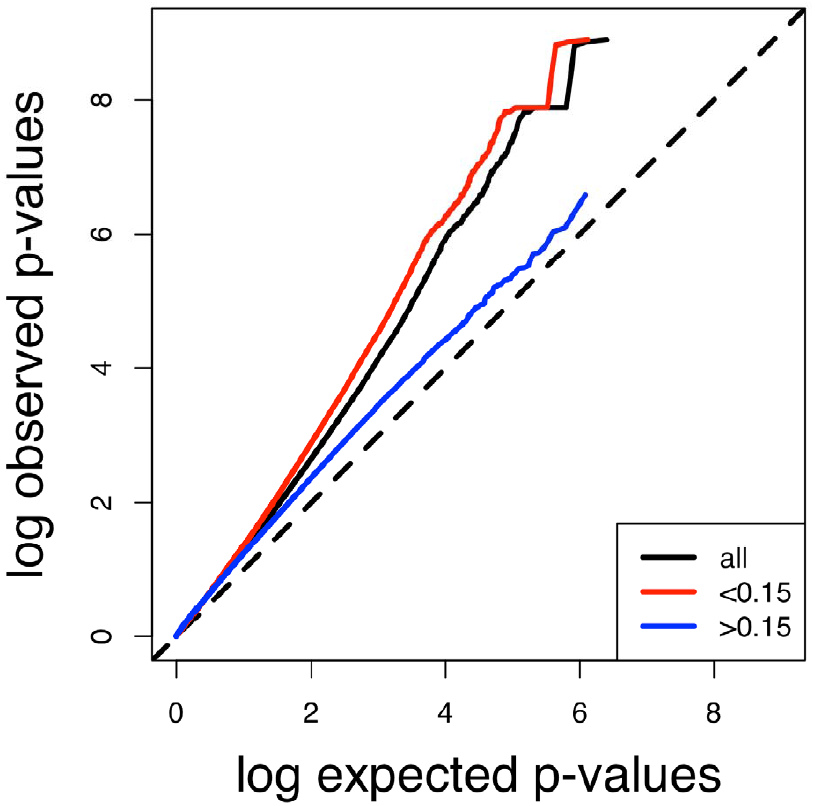
Quantile-quantile plot of observed vs. expected *P*-values genome wide. Black: all SNPs; red: SNPs with MAF < 0.15; blue: SNPs with MAF>0.15.

### Comparing multivariate and univariate analyses

To understand the relative power of the multivariate analysis, we also carried out univariate analyses of each SNP on the scores on PC1 through PC20 of the entire data set. When we applied the Storey and Tibshirani (2003) FDR algorithm independently to each of the twenty sets of P values, there were 4 significant SNPs on PC1 (shown as green dots in Figure 4) and none on the other 19 axes at a FDR of 5%. Just one of these sites is also significant at the FDR 5% level in the multivariate analysis (3L:17980378).

To further compare the multivariate and univariate results, we also applied the same critical P-value identified as the FDR 5% cutoff in the multivariate analysis (P=0.00007) to all of the univariate analyses. A total of 6,990 SNPs were identified as significant at P<0.00007 in at least one univariate analysis. Only 139 of these were also significant in the multivariate analysis. In addition, only 24 sites were identified as significant in two different univariate analyses. Figure 4B shows the genomic locations of the 565 sites significant at P=0.00007 on PC1.

To understand the nature of the differences in power between the multivariate and univariate analyses, we plotted measures of both univariate and multivariate effect size in Figure 5, classified by whether they were significant in the corresponding univariate analysis at P<0.00007. SNPs identified as significant in the univariate analyses have much larger than average effects on the PC that they are significant for (red squares) compared to the average effect of all other SNPs on that PC (blue circles). For SNPs significant on low-ranked PCs, the multivariate vectors are close to the average vector length of all SNPs. In contrast, the average score of a SNP that is significant in the multivariate analysis (green diamonds) is modestly higher than average across the full range of PCs. These comparisons suggest that the univariate analyses identify SNPs whose effects are unusually concentrated on just that PC, but are otherwise unremarkable.

**Figure 5.**
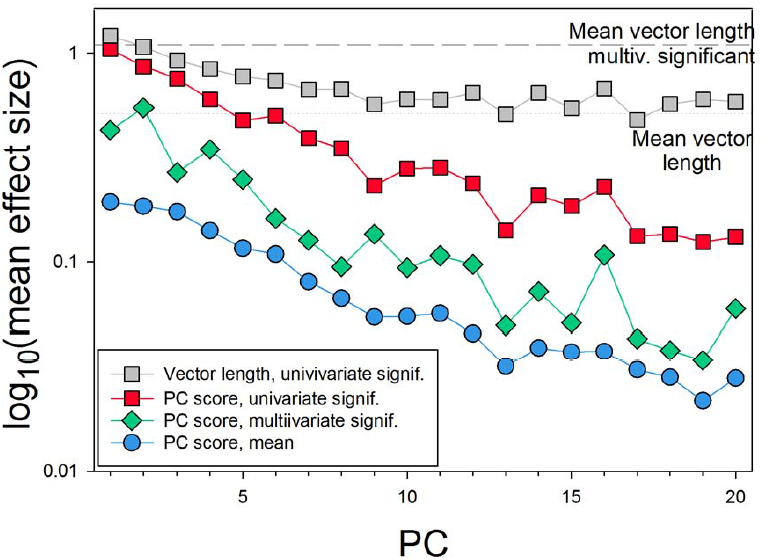
Mean measures of multivariate and univariate effect size for SNPs categorized by significance of the univariate test on each PC using P=0.00007 as a cutoff. Grey squares: total multivariate effect size for SNPs significant in the corresponding univariate analysis; Red squares: univariate effect size for SNPs significant in the corresponding univariate analysis (also shown in Figure 5B); Green diamonds: univariate effect size score for SNPs significant in the multivariate analysis; Blue circles: univariate effect size score for all SNPs. Horizontal reference lines show the mean multivariate effect size for all SNPs and for all SNPs significant in the multivariate analysis.

### Correcting for the effects of linkage disequilibrium

Despite the relative lack of population structure in the DGRP population (Mackay *et al.* 2012), there is nevertheless substantial linkage disequilibrium (LD) (Huang *et al.* 2014; Houle and Márquez 2015; Pool 2015). The average number of highly correlated SNPs (*r*^2^ ≥ 0.5) with each significant SNP is very large at low MAF, but still substantial at high MAF, as shown in Figure S2. The probability is greater than 0.5 that at least one other SNP in the genome is highly correlated with each significant SNP at all MAF, as shown in Figure S3. More striking is the fact that SNPs with low MAF have a substantial probability of being correlated with SNPs more than 100kb distant. We interpret this as being due to ‘rarity disequilibrium’ (Houle and Márquez 2015) due to the large number of low MAF SNPs, and the relatively few combinations of line genotypes that can generate a low MAF as opposed to a high MAF. Twenty-five percent of the SNPs that we analyzed have MAF<0.06, and 50% have MAF<0.137. While there is a difference in the mean number of correlated SNPs between regions inside and outside of inversions, the probability that there is at least one such correlation is affected very little by inversions (Houle and Márquez 2015).

These results suggest that the statistical signal from a focal SNP will very often be confounded with effects of other correlated SNPs, and more importantly, that those correlated SNPs will often be so distant from the focal SNP that incorrect inferences are likely to be drawn about genes implicated by a significant association. Furthermore, methods for adjusting the FDR all assume that the tests are themselves independent, which is violated for correlated SNPs. The simple MANOVA results presented above are thus likely to be misleading in many cases. We implemented several additional analyses to help judge the likelihood that a SNP with a significant test result in the MANOVA was a causal SNP, or likely to be close to a causal SNP.

A practical challenge to analyses with more predictor variables is the number of missing calls in the DGRP. The sample of lines that can be used falls rapidly when analyses are confined to genotypes at which all predictors have been called. Furthermore, imputation involves assumptions that may affect the results. In our analyses, we focused on one significant SNP at a time, and then considered the family of potential “competitor” SNPs that consists of each significant SNP that is also closest to the same gene, and all the SNPs that are highly correlated with any of these significant SNPs, including the focal SNP, plus the three common inversions. Missing data for the focal SNP were not imputed, but missing data for competitor SNPs were imputed to the common SNP.

In our first analysis, we evaluated the continued significance of each of the significant SNPs in pairwise multivariate multiple regressions with each of the possible competitor SNPs. We then counted the number of SNPs that reduced the probability that the focal SNP is significant, and determined how far they are from the focal SNP. Of the 2396 significant SNPs at FDR 5%, 96 had no other SNPs that affected the significance of the focal SNP, while an additional 249 were not affected by any SNPs mapping more than 5kb away.

In the second analysis, we carried out a stepwise multivariate multiple regression to evaluate whether the focal SNP remains significant when other predictors are included in the model. Starting with a model with just the focal SNP, we allowed up to six stepwise additions of other predictors, as described in the Methods section. Of the 2396 focal SNPs, 1577 retained significance in this analysis. These two analyses are complementary, as the first asks whether the causal signal from each SNP disappears due to “competitor” SNPs, while the second asks whether the SNP retains explanatory power in a combined analysis with other explanatory SNPs. Putting these two sets of results together, we consider the gene implicated by the SNP to be interpretable if it remains significant in the multivariate multiple regression and is not more than 5kb from the farthest SNP that causes it to lose significance. After this filtering, there were 239 SNPs which implicated small genomic regions as very likely to have effects.

Finally, we performed a cluster analysis to group SNPs according to their LD. We identified a total of 862 “clusters,” including 659 singleton clusters which correspond to significant SNPs uncorrelated (at r^2^≥0.5) with any other significant SNP. We treated these SNPs as a subset of statistically independent sites to investigate functional associations of genotypic variation, as described below. At the other extreme, two large clusters contain 236 and 644 SNPs, respectively, including correlations between both short- and long-distance (>1 Mb or in different chromosomes) SNPs.

### Appendage development implicated by GO analysis

We performed gene ontology analysis for the closest genes to the associated SNPs. The 1577 SNPs that implicated specific genomic regions were near 1188 different genes, while the 239 filtered SNPs implicated 196 genes. For both sets of genes, the GO categories anatomical development and morphogenesis (organ development and organ morphogenesis in both) and neurogenesis (generation of neurons and neuron differentiation) show evidence of being overrepresented (Figure S4) relative to the complete set of *Drosophila* protein coding genes. The genes implicated include many components of the planar cell polarity pathways known to influence final wing form (i.e. *fat*, *dachsous*, *scribbler*, *grunge*) and signaling molecules crucial to wing developmental and vein specification (i.e. *vein*, *apterous*, *Egfr*, *sulfateless*, *doc2*). The neuronal development genes over-represented include many genes expressed at the wing margin.

## Validation of SNP effects by phenotypic effects of expression knockdowns

As one validation of putative causal SNPs, we utilized quantitative knockdowns of gene expression at 97 different genes using RNAi with a gene-switch (mifepristone-dependent) tubulin-GAL4 line (see Methods; experiments listed in File S3). Figure 6A shows the effects of knockdowns at *Egfr* on wing shape at four different levels of mifepristone. To summarize these results we performed a multivariate regression of size and shape on mifepristone levels to obtain a single summary vector. The *Egfr* regression vector is shown in the left wing in Figure. 7B. We call the set of phenotypic alterations observed on knockdown a dictionary of genetic effects. We note that dictionary knockdowns reduce gene expression throughout the body during the entire duration of wing development. The effects of the knockdowns may be different from those of SNPs, even if the regions implicated in our analyses have phenotypic effects mediated by changes in gene expression,

**Figure 6.**
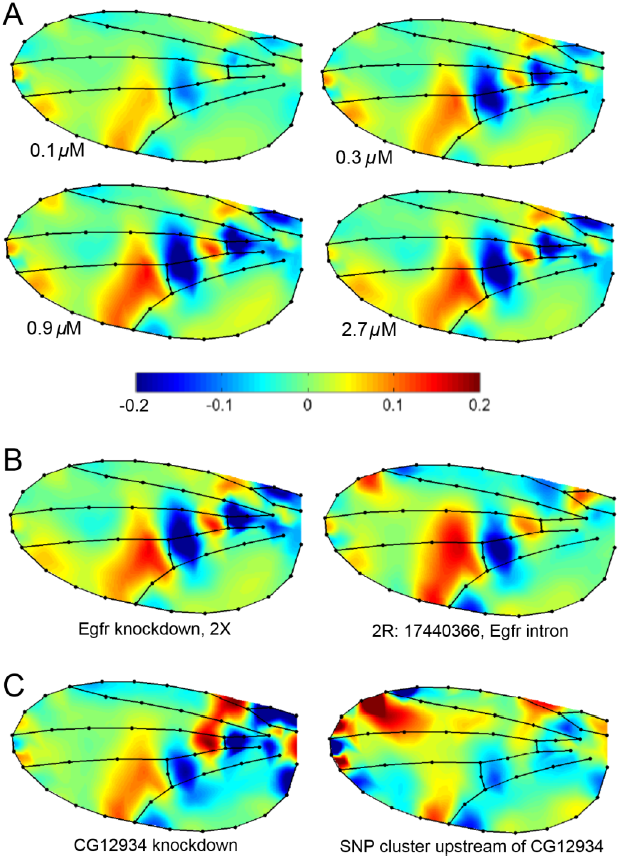
Wing shape deformations inferred for gene knockdowns and SNP effects. (A) Effects of different levels of *Egfr* knockdown on wing shape. (B) Comparison of knockdown(left) and SNP vectors (right) for *Egfr*. The *Egfr* knockdown is the regression of the shape changes shown in (A) on the level of mifepristone applied. (C) Comparison of knockdown (left) and SNP LD cluster vectors (right) for CG12934.

We compared dictionary effects to the effects of LD clusters in the DGRP. We excluded clusters containing SNPs more than 5kb apart and those with SNPs closest to the coding regions of more than one gene, which left a total of 792 LD clusters. Twenty-five of the genes in the dictionary matched the closest gene to an LD cluster containing significant SNPs, and were substantially expressed in the wing disc (Table 2). Eight of these genes matched two LD clusters, and one matched three LD clusters, giving a total of 35 dictionary genes that match interpretable LD clusters. When more than one SNP in a cluster was significant, we averaged their effects weighted by the allele frequency, *p*, as *p*(*1-p*).

We compared the directions of effects in the entire phenotype space as the absolute value of the vector correlations between cluster and dictionary effects. Vector correlations are only influenced by the direction of effects, and not by their magnitudes. Our estimates of SNP effect direction, however, have high sampling variance when the magnitude of an effect is small.

We evaluated whether these dictionary and LD cluster effects were significantly correlated at several different levels. First, we compared the entire set of correlations to see if they were higher than expected based on comparison with randomly selected vectors drawn from the analysis of the entire set of SNPs (see Methods for details). The average correlation was 0.09 units higher than expected, which was significantly different from 0 at P<0.02. This inference of correlation is reinforced because the vector lengths of both the dictionary and the SNP effect have significant Spearman correlations with their vector correlations (dictionary *r*_*s*_=0.37, P=0.026; clusters *r*_*s*_=0.34, P=0.041).

Table 2 gives the results of tests for greater than expected vector correlations at the gene and LD cluster level. We examined whether any of the genes had at least one LD cluster more highly correlated with the dictionary effect than expected, given the number of LD clusters that correspond to that gene. Two genes (*CG12934* and *Egfr*) were significantly correlated at *P*<0.05; and four had *P* values between 0.1 and 0.05. At the level of the LD cluster, there were three significant matches – one of the two *CG12934* clusters at *P*<0.01, one of two clusters at *luna*, and the single cluster at *Egfr* at *P*<0.05; an additional 7 clusters yielded *P*-values between 0.1 and 0.05. LD cluster and dictionary effects at *Egfr* and *CG12934* are shown in Figure 6B-C. These results are consistent with the finding of a bias towards high correlations, and shared directions of genotypic effects.

**Table 2.**
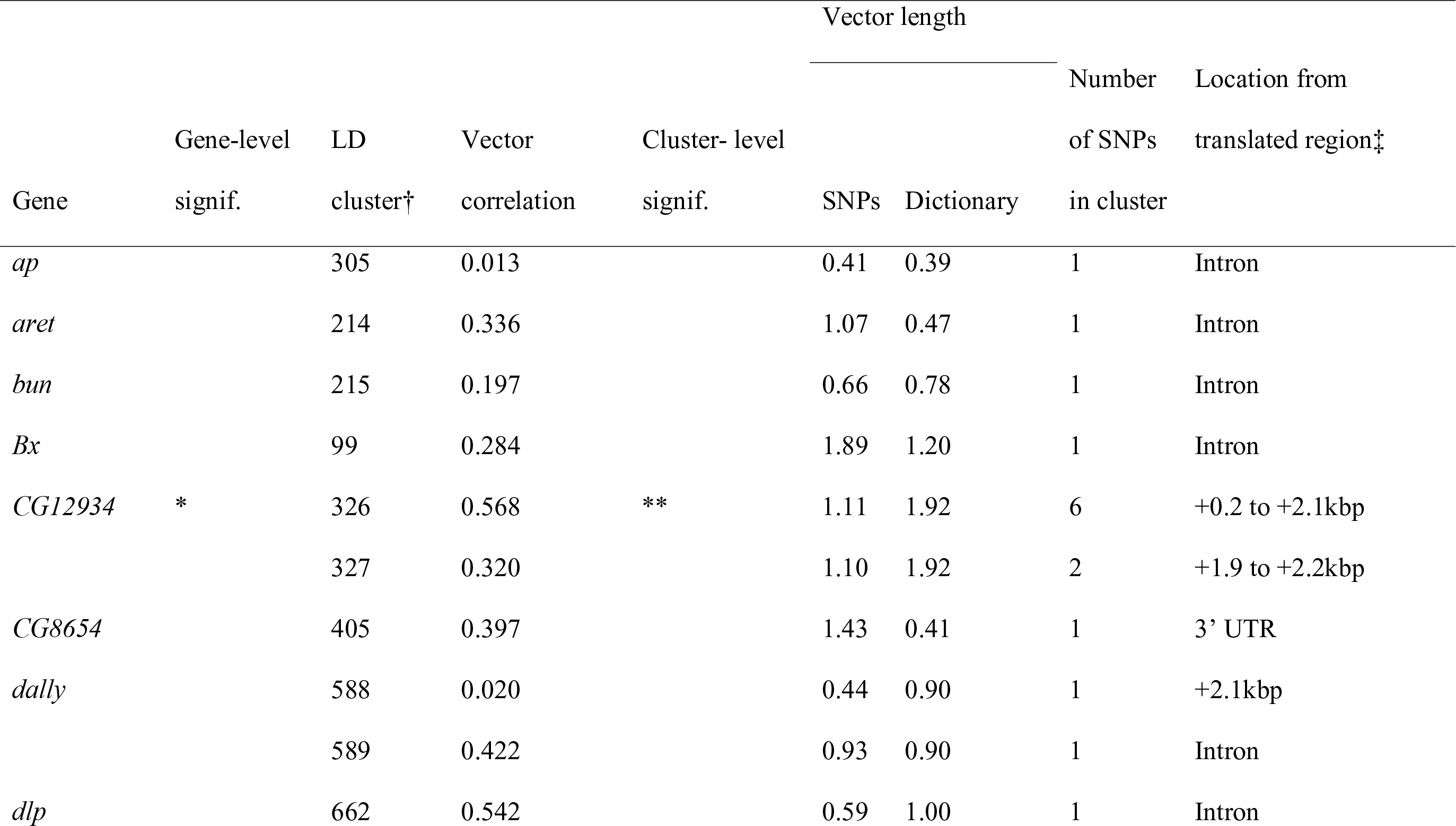
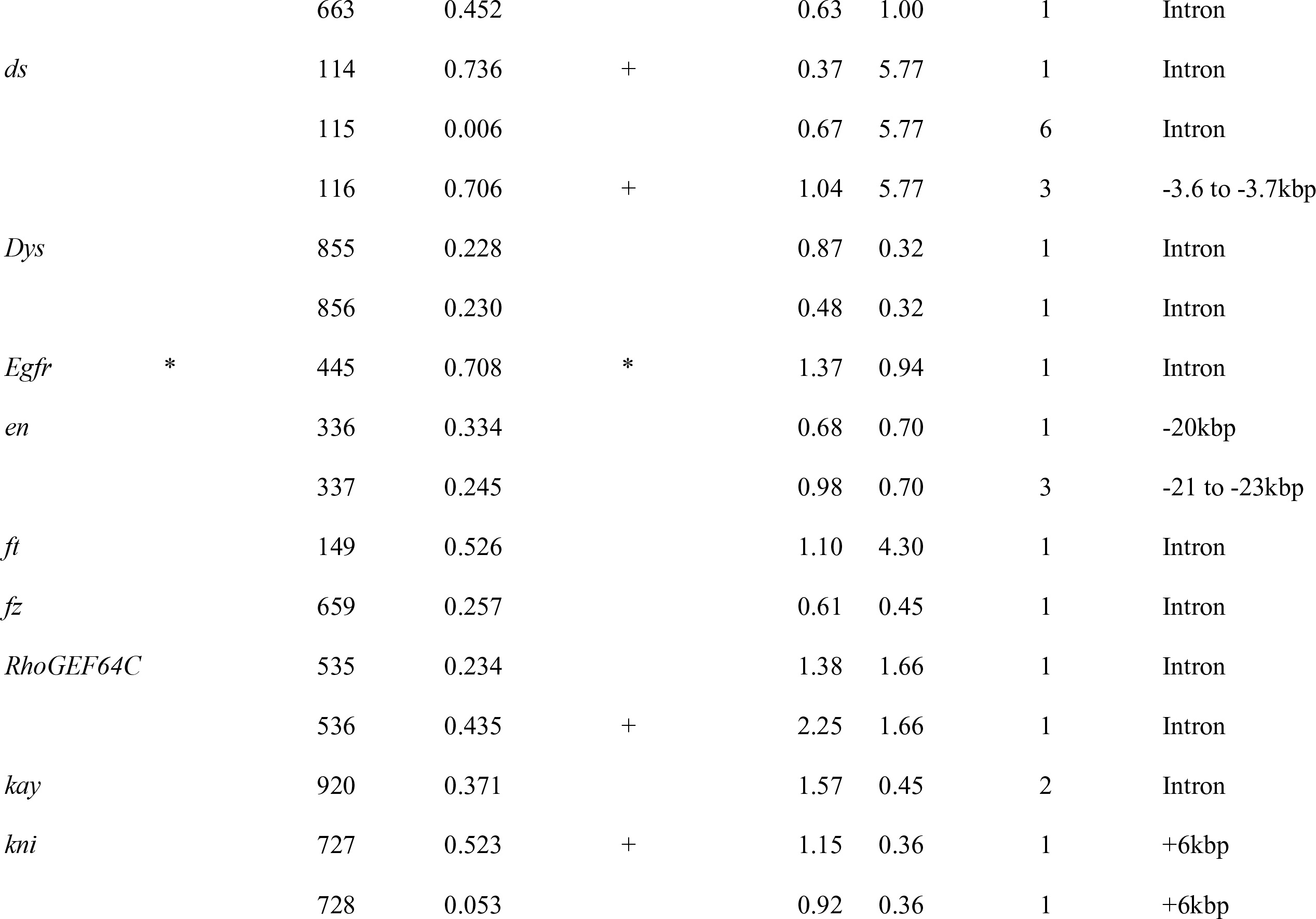
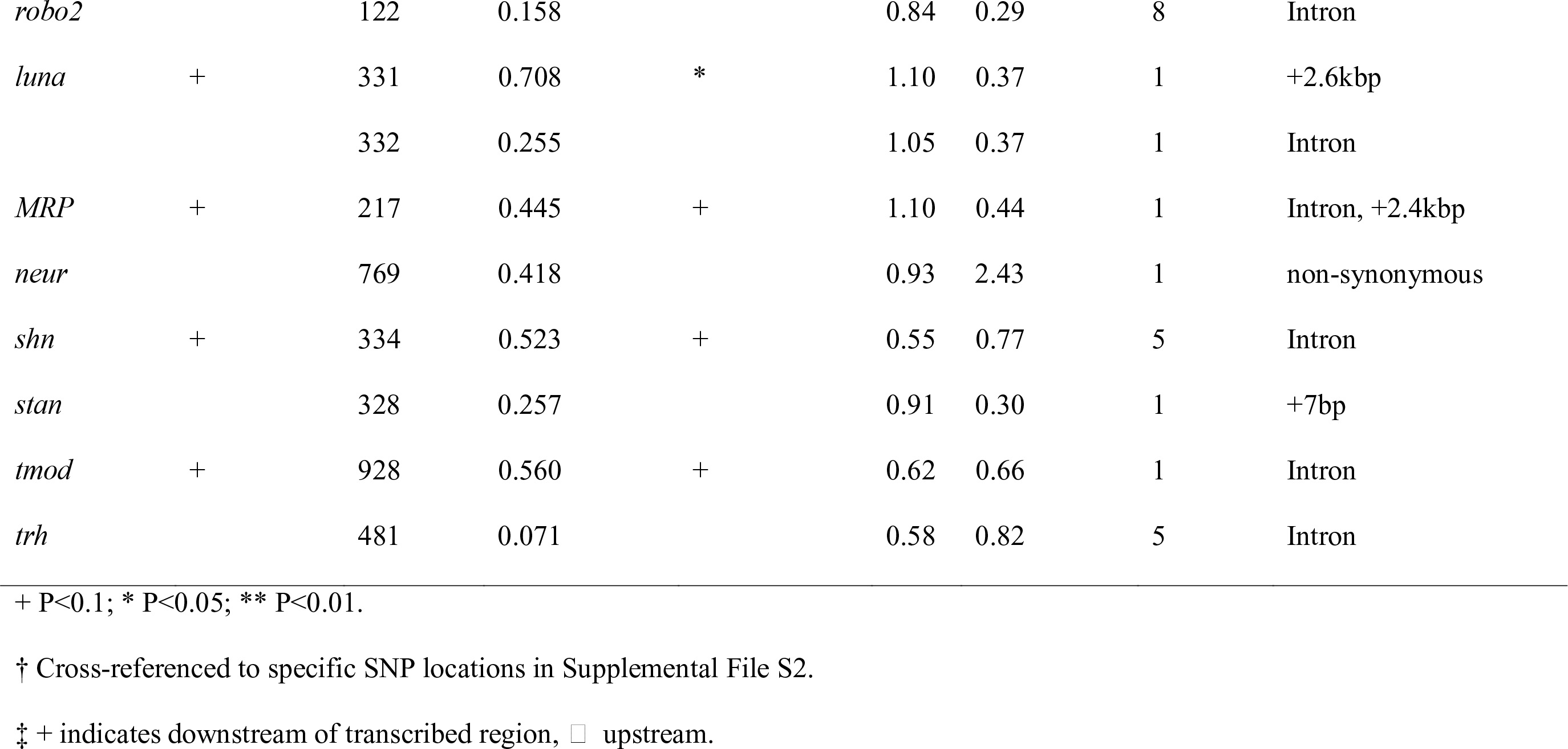
**Comparison of knockdown (Dictionary) effects on genes with significant SNPs close to the coding region.**

## Vector correlations between gene knockdowns and other SNP effects

In addition, we examined the vector correlations between each of the dictionary (gene knockdown) vectors and the effects of each of the 792 LD clusters of significant DGRP SNPs that included SNPs less than 5kb apart. Given the large number of non-independent tests, significance of the vector correlations was evaluated using a structured randomization test that controlled the knockdown-wise error rate (see methods). Vectors correlations *r*>0.5 are listed in File S4. Examination of the significant vector correlations revealed several clusters of highly correlated effects that implicate particular pathways in the production of phenotypic variation. SNPs in *ds* had significant correlations with the knockdowns in *nemo* (*r*=0.84), *ex* (*r*=0.78), *wts* (*r*=0.71) and suggestive correlations with knockdowns of *ds* itself (Figure S5). This is particularly interesting as these are in the fat-hippo pathways that influence both growth and planar cell polarity, and strongly influence final wing shape and size (Rogulja *et al.* 2008; Zecca and Struhl 2010; Schwank *et al.* 2011; Irvine 2012). We also note the significant phenotypic vector correlations between SNPs in the *Dorsocross2* gene with RNAi knockdown of the *Ultrabithorax* gene (*r*=0.87), which reflects a recently identified set of functional associations with respect to wing and haltere development (Sui *et al.* 2012; Ibrahim *et al.* 2013; Simon and Guerrero 2015). In addition, there are numerous other associations with SNPs in genes not currently annotated for their influence on wing development.

## Validation in the Maine and NC populations

Of the 321 SNPs that we had selected for validation testing from our GWAS based on DGRP Freeze 1 genotype data, 284 were found with a high enough MAF in Freeze 2 to enable validations of the association analysis of the DGRP results. Of these, only 49 were still highly implicated in our multiple multivariate regression results using the freeze 2 genotype data. We first investigated whether, as a set, these SNPs chosen for validation showed a greater similarity of direction of SNP effects across the two data sets compared with 1000 random partitions of the data (with 284 SNPs in each partition). Over the random 1000 subsets, the average vector correlation was ~0.224. For the full set of 284 SNPs chosen for validation, the vector correlation was ~0.23, but was not outside of the range of expected values based on the random subsets. However, when we only examine the subset of 49 SNPs which showed some evidence of association in our final analysis, the vector correlation increased (~0.24) and exceeded the 95% intervals. Similarly, when we investigated the correlation among effect sizes for the significant DGRP SNPs with the validation set, the Pearson correlation was 0.67, and exceeded the correlations produced with all of the 1000 random subsets.

Despite, this, at the level of individual SNP validations, we observed little evidence of replication, with 28 of the SNPs showing validation in the MENC data set, with an alpha set at a nominal value of 0.05, and only 3 with P-values below 0.001.

## Discussion

The results of our fully multivariate genome-wide association analysis have implications for the study of inheritance and evolution of the *Drosophila melanogaster* wing, for the genetic architecture of quantitative traits, for the study of the genotype-phenotype map and for the usefulness of multivariate association analyses. We discuss each of these in turn.

### Inheritance of Drosophila wing shape

The *Drosophila* wing is a single structure, consisting of veins connected by wing blade tissue. The integration enforced by the physical connection between each part of the wing, and the continuity of these structures throughout development makes it a natural subject for a multivariate genetic analysis. Any change during development that affects one aspect of the wing, such as the length of a particular vein, must also affect adjacent areas of the wing. The processes most likely to affect wing shape and size are the pattern of growth of the wing tissue, the differentiation of veins from non-vein tissue, and the rearrangement and planar polarization of cells during pupariation (Matamoro-Vidal *et al.* 2015). The known candidate genetic pathways that affect these key developmental events have effects across broad regions of the wing, rather than being confined to one small area. These considerations suggest that it is impossible to choose a genetically independent set of wing traits to measure.

A second layer of dependency among measurements of the wing is introduced by the geometric morphometric analysis we used (Zelditch *et al.* 2004). There is no one reference structure on a complex integrated morphological structure like a wing that can be used as a standard to compare with the locations of other structures. One can only interpret the relative locations of all measured structures to one another, giving one more reason why it will not be possible to define a set of traits that can be measured independently. Both of these features suggest that pleiotropy is an absolutely unavoidable feature of variation in continuous morphological structures, such as the fly wing. Consequently, the pattern of effects on all measured phenotypes in the wing will be more useful than any subset of measured variables.

The multivariate analysis of variance (MANOVA) that we used in this study can be thought of as consisting of two steps – first, identifying the direction in phenotype space that best differentiates the two genotypes at a particular genomic site, and second determining whether the magnitude of the difference in that direction is sufficiently large to warrant our attention. The first step defines a trait – the direction in phenotype space that maximizes the distinctness of the means of the two genotypes. Second, statistical significance is estimated based on how unlikely that difference is under the null hypothesis. In the more typical series of univariate analysis, traits are chosen from a finite set of possible measurements that cannot capture the entirety of phenotypic effects, except in the unrealistic case of uncorrelated traits. The multivariate analysis uses the data to decide which of the infinite combinations of the finite measurements best shows the difference between genotypes. This is why multivariate analyses are more powerful than univariate ones.

In addition, this aspect of MANOVA corresponds to our intuition about what variant genotypes that actually affect the integrated wing phenotype should do: in principle, every site affecting wing development could do so in a slightly different way, and each of those changes will have pleiotropic effects that extend across the wing. The plots of wing shape change in Figure 6B-C and Figure S5 represent estimates of those directions of some of our significant SNPs.

For our data, the gain in power in the multivariate analysis was very dramatic. At a false-discovery rate of 5%, 2,396 SNPs were identified as potentially significant in the fully multivariate analysis. In comparison the univariate analyses of principal component (PC) scores identified just four significant variants on PC1, and none on the next 19 PCs when using the same FDR algorithm on each axis. When we utilized the P-value estimated from the FDR from the multivariate analysis (P=0.00007), almost 7,000 SNPs were nominally significant, but just 24 of those had significant effects on two or more PC axes. It is particularly notable that just 139 SNPs had significant effects in both the multivariate and at least one univariate analysis. We interpret this to mean that the false discovery rate of the more liberal (P=0.00007) univariate criterion is quite high. The univariate analyses detect sites with effects that are unusually concentrated along one PC axis, but look unremarkable in the entire phenotype space.

Previous association studies on aspects of wing shape in *D. melanogaster* have also detected relatively large numbers of QTLs, given the number of markers employed. Weber *et al.* (1999; 2001) generated recombinant inbred lines (RILs) between populations selected for high- and low-values of a univariate wing shape index, and found at least 20 sites over the two largest chromosomes with uniformly small effects. Zimmerman *et al.* (2000) found evidence for a dozen QTLs for several aspects of wing shape in each of two small mapping populations, each founded by crossing two inbred lines. Mezey *et al.* (2005) mapped at least 21 QTLs for the first seven principal components of wing shape in a set of RILs derived from the cross of a single pair of wild-collected flies.

The large numbers of sites implicated in both the present and previous studies, strongly suggests that the inheritance of wing size and shape is highly polygenic, with many genetic variants of small phenotypic effect segregating in natural populations of *D. melanogaster*. In this study, effect sizes are relatively uniform and no large effects were detected; the median proportion of variance explained by a statistically significant SNP, averaged within LD cluster, is 1.1%, while the maximum is just 3.6%. This pattern is reminiscent of the genetic architecture of human height, where a large number of sites with individually small effects are responsible for the standing variation (Lango Allen *et al.* 2010; Yang *et al.* 2010; Wood *et al.* 2014). The magnitude of the effect sizes is certainly overestimated, as is the proportion of variance explained. There are two well-known causes for this overestimation that our analysis does not correct for. First, we report effect sizes at sites that are individually statistically significant. This will enrich for sites where effects are overestimated, rather than underestimated, causing the Beavis effect (Beavis 1994; Beavis 1998; Xu 2003). Second, there is substantial linkage disequilibrium involving many of our significant sites, and our analyses and, in particular, the false discovery rate calculations assume that all sites are independent. It is likely that some of our LD clusters may contain multiple causal sites, and their effects become confounded in the estimates of effect size.

We have carried out an extensive series of simulations of data sets similar to ours (Márquez and Houle 2015) that show the power of our experiments is quite modest for sites that explain just 1% of the variation, as our median significant sites are estimated to do – perhaps just 20%. Such low power means that we will detect only a minority of all the variants with an effect on the phenotype. This ensures a substantial Beavis effect. It also suggests that correlations among predictors could inflate the false discovery rate above the nominal 5% rate that we strove to achieve. Conversely, if many sites with small effects are in fact responsible for the genetic variation in wing shape, the modest departures from the null distribution of P-values revealed in the qq-plot in Figure 3 are expected. These considerations suggest that we can have considerable confidence in the overall genetic signal, but low confidence in individual sites.

Our estimates of effects on all shape traits simultaneously allows us to undertake validation experiments that test whether validation effects are in directions more similar using the angle between the observed vectors, and not just whether phenotypic effects can be detected at one trait. We performed two such sets of validation experiments, and their results are both consistent with the highly polygenic architecture with small effect sizes. The phenotypic effects of knockdowns of genes implicated in the initial GWAS, the dictionary experiment, provided good evidence that SNP effects are more similar to these than expected under the null hypothesis of no similarity. In several cases the effects of particular SNP-knockdown pairs are individually more similar than random vectors (Figure 6).

Validation of SNPs in a second panel of lines from natural populations from Maine and North Carolina (ME-NC) again suggested that overall the direction of effects is more similar to random SNP subsets than expected by chance. The evidence for validation for individual DGRP SNPs was relatively poor, and no individual sites were strongly validated. The precision with which phenotypic effects are estimated is positively correlated with effect sizes. This suggests that the direction of the many small effects we detected is imprecisely estimated, which will tend to increase the angles between effects. This effect should be particularly large comparing both the DGRP and ME-NC vectors, which are both small in magnitude, and likely overestimated in the DGRP analysis, as discussed above. In contrast, the directions of the dictionary knockdowns are larger and more precisely estimated. This may account for the stronger validation in the dictionary experiment.

Despite the difficulties in identifying particular candidate SNPs, the additional filtering step we performed using the multivariate multiple regression with “competitor” SNPs in LD with the focal SNP helped us identify a smaller set of genes more likely to be responsible for genetic variation. Indeed, both the full set of 1188 genes implicated by the full set of significant SNPs, and the filtered set of 196 genes implicated by the SNPs that showed evidence of over-representation of *Drosophila* limb development genes (Figure S2). Furthermore the high vector correlations of dictionary effects with SNP effects for individual genes such as *Egfr* (Figure 6) suggest that meaningful candidates can be identified for further verification and study. *Egfr* has previously been implicated in QTL and candidate gene association mapping of *Drosophila* wing shape (Palsson and Gibson 2000; Zimmerman *et al.* 2000; Palsson and Gibson 2004; Palsson *et al.* 2004; Dworkin *et al.* 2005). Other genes such as *dachsous* (*ds* Figure S5), have profound effects on wing morphology including overall shape, and remain important candidates for future work.

While we have demonstrated the power and prospect of a fully phenomic approach to association studies, there are important caveats to consider, as with all GWAS studies. Population sub-structure and genetic relatedness among samples can confound the independent estimation of genetic effects (Ziv and Burchard 2003; Tian *et al.* 2008). Indeed, while the DGRP inbred lines have generally been considered to have little sub-structure, recent evidence suggests that there has been some recent admixture with African populations (likely via the Caribbean) (Duchen *et al.* 2013; Pool 2015). While LD due to spatial proximity between sites has been recognized as an important concern for decades, the large number of rare alleles in natural populations gives rise to random linkage disequilibrium in the sample, termed “rarity disequilibrium” (Houle and Márquez 2015). We considered and in part controlled for this by using a second round of model fitting that includes sites in LD (both spatially proximal and distant) as covariates. Recent analytical advances, such as the use of mixed models (Yang *et al.* 2010) or regularization schemes (Peng *et al.* 2010; Wang *et al.* 2015), that minimize the overfitting engendered by simultaneous consideration of huge numbers of predictors are promising solutions to such problems. Unfortunately, their application to multivariate problems is not yet mature.

Overall, we are confident that our list of significant SNPs is enriched for causal QTNs and sites correlated with QTNs, but remain uncertain of which sites are actually causing the genetic variance we observe.

## Why multivariate association studies?

The larger point that we wish to emphasize is that multivariate analyses increase both the power of association studies, and the interpretability of the results obtained over a series of univariate analyses.

As discussed above, the power of our multivariate analyses is far greater than those of a comparable set of univariate analyses; we detected 2,396 significant SNPs at an FDR of 5% in the multivariate analyses, but just four in univariate analyses of scores on the 20 most variable principal component axes. The fact that this result has repeatedly been demonstrated in simulation studies using a variety of statistical methodologies (O’Reilly *et al.* 2012; Stephens 2013; van der Sluis *et al.* 2013; Zhou and Stephens 2014; Márquez and Houle 2015) suggests that the expectation of increased power is general. Except in special cases, any multivariate analysis will be more powerful than the corresponding set of univariate analyses.

A second important reason for multivariate analyses is that the multivariate effect vector estimated is far more informative than a series of decisions about which traits are affected by each SNP that results from standard univariate testing. We exploited this in our analyses to demonstrate that a series of gene knockdowns have effects that are more similar to our effects than expected under a random model, to pick out a few similarities that are particularly significant and deserving of further study. Highly correlated effects suggest the potential for some shared biological function.

A final critical justification for transitioning from univariate to multivariate association studies is to enable the study of the genotype-phenotype map, how genomic variation is translated into phenotypic variation (Houle 2010; Houle *et al.* 2010). Everything that we know about genetics and biology suggests that genomic variation will have pleiotropic effects. We can’t begin to study pleiotropy without studying multiple traits. Every phenotypic effect will have a molecular origin, for example in gene expression, which then ramifies outwards to cells, tissues and finally to the outward aspects of organismal form and function such as morphology and behavior. Each such molecular change may have effects on many whole organism phenotypes. For example, the study of even the simplest monogenic human genetic diseases, such as sickle-cell anemia, inevitably reveals a host of disorders tracing back to the single genetic cause. Decisions about how to treat genetic disease, the value of a genetic variant in plant or animal breeding, or whether an endangered population is likely to adapt to a changing environment will be improved when we have information about all of the pleiotropic effects of genetic variation, and not just the few that happen to have been studied.

The prevailing approach to the study of pleiotropy is to perform a series of univariate analyses to count the numbers of traits that are significantly affected by a SNP. This biases the GWAS results towards discovering just a few large effects, even if the underlying architecture is highly pleiotropic, because most GWAS have modest power (Beavis 1998; Manolio *et al.* 2009; Slate 2013). Our univariate analyses could be used to argue that there is low pleiotropy as only PC1 was affected, and no SNP had a significant effect on two PC scores, while our multivariate analyses reveal that the directions of significant effects are quite variable (Figure 8). A striking example of the bias this causes is a recent GWAS of brain regions in mice that concluded that almost all QTLs affected only a single brain region, implying no constraints on the evolution of brain shape (Hager *et al.* 2012). The results were based on a panel of 100 recombinant inbred lines, so that the power to detect effects was small. Not a single pleiotropic effect was detected based on the failure of any QTL to reach statistical significance for more than one brain region. In reality, we know that body size is affected by hundreds of segregating variants in large populations (Wood *et al.* 2014), and some of these should also affect brain size as a whole (Lande 1979). This methodology is unfortunately ubiquitous in studies of pleiotropy. A rigorous testing approach is appropriate to the goal of finding the candidate genes for further investigation, but not to the goal of estimating the pattern of pleiotropy. In the univariate case the use of statistical testing creates the problem of ‘missing heritability’ (Yang *et al.* 2010). In the multi-trait case, univariate testing introduces ‘missing pleiotropy.’

One popular alternative to a fully multivariate approach is to apply dimension-reduction techniques that redefine traits as combinations of the measured traits, such as principal components analysis and linear discriminant analysis, then analyze a small number of these linear combinations using univariate statistics (e.g., Zimmerman *et al.* 2000; Mezey *et al.* 2005). The main justification for this approach is to ensure that the traits analyzed are independent from each other. While this does provide valuable information, such analyses are still a series of univariate analyses that will be less powerful than a fully multivariate analysis, as discussed above. If, in addition, the variables analyzed do not capture all the phenotypic variation, some of the information in the original sample is not utilized.

## Dimensionality as a blessing rather than a curse

With all these advantages to multivariate association analyses, why are they still rare? In some cases, there are substantial statistical barriers to a fully multivariate analysis. For example, it is challenging to combine binomial and normal variates in the same analysis, although solutions have been proposed (e.g., O’Reilly *et al.* 2012). Many multivariate data sets have incomplete phenotypic data, and restricting the analysis to just those individuals with complete data may reduce sample size too much for reasonable inference. Multivariate methods are unfamiliar to many researchers, posing a relatively simple hurdle to their adoption.

We suspect that a final factor interfering with the widespread adoption of multivariate methods is summed up in the phrase “curse of dimensionality.” This phrase was originally coined by Richard Bellman (1957) and has since become a meme useful for causing unease about multivariate analyses, even when the nature of the curse remains implicit. It generally denotes the notion that the hypervolume of sample space increases rapidly with the number of dimensions measured, while the sample size remains fixed, resulting in data that is ever sparser as dimension increases. Zimek *et al.* (2012) identify eight separate challenges that increase with dimensionality of the data set just in the realm of distance-based analyses (such as detecting neighbors, hubs, outliers, etc.). They also note that many of these are problematic only in the limiting case where all variables are independently and identically distributed; biological data are always correlated and often clustered. Our argument that the relationship between vectors of effects is more informative in a high dimensional data space is essentially the flip side of the standard sparsity argument. Effects become more informative because a finite set of real effects must be sparser in a larger space, and therefore both similarities and differences become more informative.

Another challenge frequently posited is that a large proportion of the measurements in a high dimensional data set may be irrelevant. Indeed, our simulations show that power of an association study declines when traits without any genetic basis are measured (Márquez and Houle 2015). Given that the current standard approach to GWAS includes just a few traits, we are confident that the number of traits can usually be greatly increased without reaching this limit. Biological measurements are expensive and time-consuming to make, ensuring that considerable thought will be expended on what to measure. Furthermore, the appropriate dimension for analysis can be estimated from data on related individuals (Kirkpatrick and Meyer 2004; Meyer and Kirkpatrick 2005; Meyer and Kirkpatrick 2008). In general, principal components analysis can reveal how much new information is added when another trait is measured, and a cutoff that seems likely to capture most genetic variation chosen.

The best answer to the concern that dimensionality can be a curse are analyses of simulated data sets that show that the power of multivariate analyses is higher, sometimes much higher, than univariate analyses. It is especially notable that, many independent simulation studies that make different assumptions, and apply a wide variety of well-established or experimental multivariate analyses all obtain this result. Most of these studies also analyze real data sets and invariably find more associations in multivariate than univariate analyses (O’Reilly *et al.* 2012; Stephens 2013; Scutari *et al.* 2014; Zhou and Stephens 2014). Our results are consistent with this pattern.

We believe that researchers should invoke the blessings of dimensionality, rather than its potential to be a curse. Multivariate analyses will generally be more powerful. The ability to estimate the direction of effects becomes more salient with the dimension of the space studied. The phenomenon of pleiotropy simply cannot be studied unless multiple traits are studied together. The prevailing estimates of pleiotropy based on sequential univariate analyses will greatly underestimate the degree of pleiotropy. Those interested in the inheritance of complex traits and the genotype-phenotype map should adopt multivariate approaches whenever it is feasible to do so.

## Data availability

Data on which this study is based will be archived in Dryad doi:XX.xxxx/dryad.xxxxxx).

## Supplemental Information

Submitted separately.

**Figure S1. Inter and intra-lab repeatability:** A) Vector correlations of deviations of line means from the grand mean. The vast majority of estimates are highly correlated. Estimates with large angles are those with small effect sizes. B) Estimates of wing size in experimental replicates in the Dworkin lab.

**Figure S2. Mean and median number SNPs correlated at r**^**2**^**>0.5 with significant SNPs, as a function of distance between sites and MAF.**

**Figure S3. Probability that a significant SNP is correlated at r**^
**2**^**>0.5 with at least one other site in the genome as a function of MAF.**

**Figure S4. Gene Ontology analysis of LD-filtered genes from Web Gestalt.**

**Figure S5. Effects of variants near the dachsous (*ds*) gene and knockdown vectors significantly correlated with them.** Two LD clusters of SNPs near *ds* contained SNPs with significant phenotypic effects, cluster 114 and cluster 116. These vectors are themselves correlated at r=0.67, and have suggestive correlations with the *ds* knockdown phenotype (lower left; r=0.75. P<0.1 LD114; r=0.70, ns LD116), suggesting a general effect of *ds* expression. The effect of cluster 114 is significantly correlated with knockdown of *expanded* (*ex*) (r=0.78, P<0.01; r=0.71, P<0.1 with LD116). The effect of cluster 116 is significantly correlated withknockdowns of *nemo* (r=0.84, P<0.001; r=0.63, ns with LD114) and *warts* (*wts*; r=0.71, P<0.001; r=0.49, ns with LD114).

**File S1. Means and S.D. of PC scores by lab and sex of fly, eigenvectors of the combined data set, and the estimated among-line genetic variance-covariance matrix in a 40-dimensional and phenotypic (97-dimensional) spaces.**

**File S2. Significant SNPs from the multivariate analysis and the results of two analyses to determine whether the SNP is likely to have a causal effect.**

**File S3. List of RNAi knockdown experiments, the quantiles of correlations under the null hypothesis of no relationship, and the angles between effect vectors and the first 5 PCS of the combined data set.**

**File S4. Vector correlations with r>0.5 between knockdown experiments and average effects of LD clusters.** Only LD clusters with sites <5kb apart and whose members are all closest to a single gene are included.

## Acknowledgements

We thank Rosa Moscarella for managing the RNAi knockdown experiments, and the many undergraduates who aided in rearing flies and imaging and splining of wings. Computing support was provided by iCER and the High-performance computing center at MSU, and Research Computing Center at FSU. We thank Trudy Mackay and the Mackay lab for providing the DGRP lines, the Bloomington Stock Center for providing stocks, and High Performance Computing for computing resources at North Carolina State University. This work was supported by National Institute of General Medical Sciences 1R01GM094424-01 to ID & DH, and U.S. National Science Foundation Division of Environmental Biology (www.nsf.gov) 0129219 to DH and EJM. The funders had no role in study design, data collection and analysis, decision to publish, or preparation of the manuscript.

